# The nitrogen-fixing symbiotic cyanobacterium, *Nostoc punctiforme* can regulate plant programmed cell death

**DOI:** 10.1101/2020.08.13.249318

**Authors:** Samuel P. Belton, Paul F. McCabe, Carl K. Y. Ng

**Author notes:** Author contributions S.P.B., P.F.M, and C.K.Y.N conceived of the study and designed the experiments. S.B. performed all the experiments and analysed the data. S.B. prepared the draft of the manuscript with the help of P.F.M and C.K.Y.N. All authors read, edited, and approved the manuscript.

## Abstract

Cyanobacteria such as *Nostoc* spp. can form nitrogen-fixing symbioses with a broad range of plant species. Unlike other plant-bacteria symbioses, little is understood about the immunological and developmental signalling events induced by *Nostoc* cyanobionts (symbiotic cyanobacteria). Here, we used suspension cell cultures to elucidate the early molecular mechanisms underpinning the association between cyanobionts and plants by studying the effects of conditioned medium (CM) from *Nostoc punctiforme* cultures on plant programmed cell death (PCD), a typical immune response activated during incompatible interactions. We showed that *N. punctiforme*-CM could suppress PCD induced by a temperature stress. Interestingly, this was preceded by significant transcriptional reprogramming, as evidenced by the differential regulation of a network of defence-associated genes, as well as genes implicated in regulating cell growth and differentiation. This work is the first to show that cyanobionts can regulate PCD in plants and provides a valuable transcriptome resource for the early immunological and developmental signalling events elicited by *Nostoc* cyanobionts.

## Introduction

Species of the heterocystous cyanobacteria genus *Nostoc* can form nitrogen-fixing symbioses with several vascular and non-vascular plants (Adams et al., 2012). Unlike other nitrogen-fixing symbioses, such as the associations formed between Rhizobia and legumes or between *Frankia* spp. and actinorhizal plants, the molecular dialogue which takes place between *Nostoc* spp. and their various host plants are less well understood (Cissoko et al., 2-18; Geurts et al., 2016; Mus et al. 2016;). Most of the cross-species signalling events which have been characterised to date primarily relate to those surrounding the exchange of macronutrients and the regulation of cellular differentiation in the *Nostoc* cyanobiont(Adams et al., 2012; Ekman et al., 2012). However, there has been little focus on the immunological and developmental signalling events which are elicited in the host to facilitate colonisation.

Incompatible plant-bacteria interactions are characterised by the perception of microbe-associated molecular patterns (MAMPs) via host pattern recognition receptors (PRRs). This rapidly induces the transcription of suites of defence-associated genes which confer pattern-triggered immunity (PTI) (Saijo et al., 2018). Similar to symbiotic associations between Rhizobia and legumes, and *Frankia* and actinorhizal plants, *Nostoc*-plant symbioses represent compatible plant-bacteria interactions. Rhizobia produce lipochitooligosaccharides (LCOs) termed Nod factors that are induced by flavonoids derived from prospective plant hosts (Mus et al., 2016). These Nod factors and putatively, exopolysaccharides, are capable of suppressing MAMP-induced defence responses (Liang et al., 2013; Wang et al.,2018).

*Nostoc* spp. possess lipopolysaccharides (LPOs) in their outer membranes and peptidoglycans in their cell walls, both of which are typical MAMPs (Erbs and Newman, 2012). However, they do not appear to possess a LCO biosynthetic pathway, but like Rhizobia and *Frankia* spp., are able to form intimate and in some cases, intracellular associations without eliciting PTI (Geurts et al., 2016). It is not entirely clear whether *Nostoc* LPOs are simply unrecognisable as MAMPs by plant PRRs, or whether symbiotic *Nostoc* spp. can actively suppress plant immune responses, such as programmed cell death (PCD). In part, the lack of attention to this aspect of *Nostoc*-plant symbioses can be attributed to the lack of host genome sequences, although this may now change as a number have recently been reported (Li et al., 2018; Pederson et al., 2019).

Originally isolated from the coralloid roots of a cycad, *Macrozamia* spp. (Rippka et all, 1979), *Nostoc punctiforme* strain ATCC 29133 (PCC 73102) is a model species of cyanobiont that has been shown to successfully colonise a broad range of plant hosts, including the hornwort *Anthoceros punctatus* (Ekman et al., 2013), herbaceous angiosperms of the genus *Gunnera* (Chiu et al., 2005), and more recently, rice (Álvarez et al., 2020). Interestingly, symbiotically competent *Nostoc* strains have also been shown to exhibit chemotaxis towards non-host plants like *Trifolium repens* (white clover) and the model dicotyledon, *Arabidopsis thaliana* (Nilsson et al., 2006). Here, *A. thaliana* suspension cell cultures were used as a model system to investigate whether *N. punctiforme* can regulate plant PCD. Additionally, to better understand the molecular dialogue between *N. punctiforme* and plants, the transcriptome of *A. thaliana* cells treated with *N. punctiforme*-conditioned medium (CM) was sequenced and analysed. Our data provide novel insights into the early immunological and developmental signalling events elicited by the *Nostoc* cyanobiont.

## Results

### *N. punctiforme* can suppress plant programmed cell death (PCD)

To investigate whether *N. punctiforme* can regulate PCD, we used a well-established method involving a 10-min, 51°C temperature stress to induce PCD in *A. thaliana* suspension cell cultures (Kacprzyk et al., 2017) following a 1-hr pre-incubation with *N. punctiforme*-conditioned medium (CM; 20% v/v). This workflow is depicted in Fig. 1a. As a control treatment, cells were pre-incubated with fresh *N. punctiforme* growth medium (20% v/v). Twenty-four hr after applying the temperature stress, cell viability in cultures pre-incubated with fresh growth medium was reduced to ∼68%, with ∼30% of cells committing to PCD, and <0.25% of cells dying by necrosis (Fig. 1b). However, when cells were pre-incubated with *N. punctiforme*-CM, viability was maintained at a significantly higher level (∼85%) whereas PCD levels were significantly attenuated (∼14%) compared to control cells (*P* ≤ 0.001; Fig. 1b). Importantly, necrosis levels did not change (Fig. 1b), which suggests that *N. punctiforme*-CM could specifically suppress PCD.

### *N. punctiforme*-CM elicits major transcriptomic changes

As a 1-hr pre-incubation with *N. punctiforme*-CM sufficed to suppress PCD, we hypothesised that a differential induction of stress-associated genes would be discernible at this time-point. To investigate this, RNA was extracted and a cDNA library generated ahead of paired-end RNA-sequencing. After applying a 0.3 FPKM cut-off, a total of 19,064 genes were determined to be expressed, 396 of which were unique to *N. punctiforme*-CM treated cells (Fig. 2a). We observed that between control and CM-treated cells, 7,032 genes were differentially expressed after controlling for false discovery (*P*adj ≤ 0.05; Supplementary Data 1). Hierarchical clustering analysis revealed that large numbers of these genes grouped into similar patterns of expression between treatments (Fig. 2b). After applying a threshold of a 4-fold change in expression, we deemed 483 and 479 genes to be significantly up- and down-regulated, respectively in response to *N. punctiforme*-CM (Fig. 2c).

**Fig. 2.**
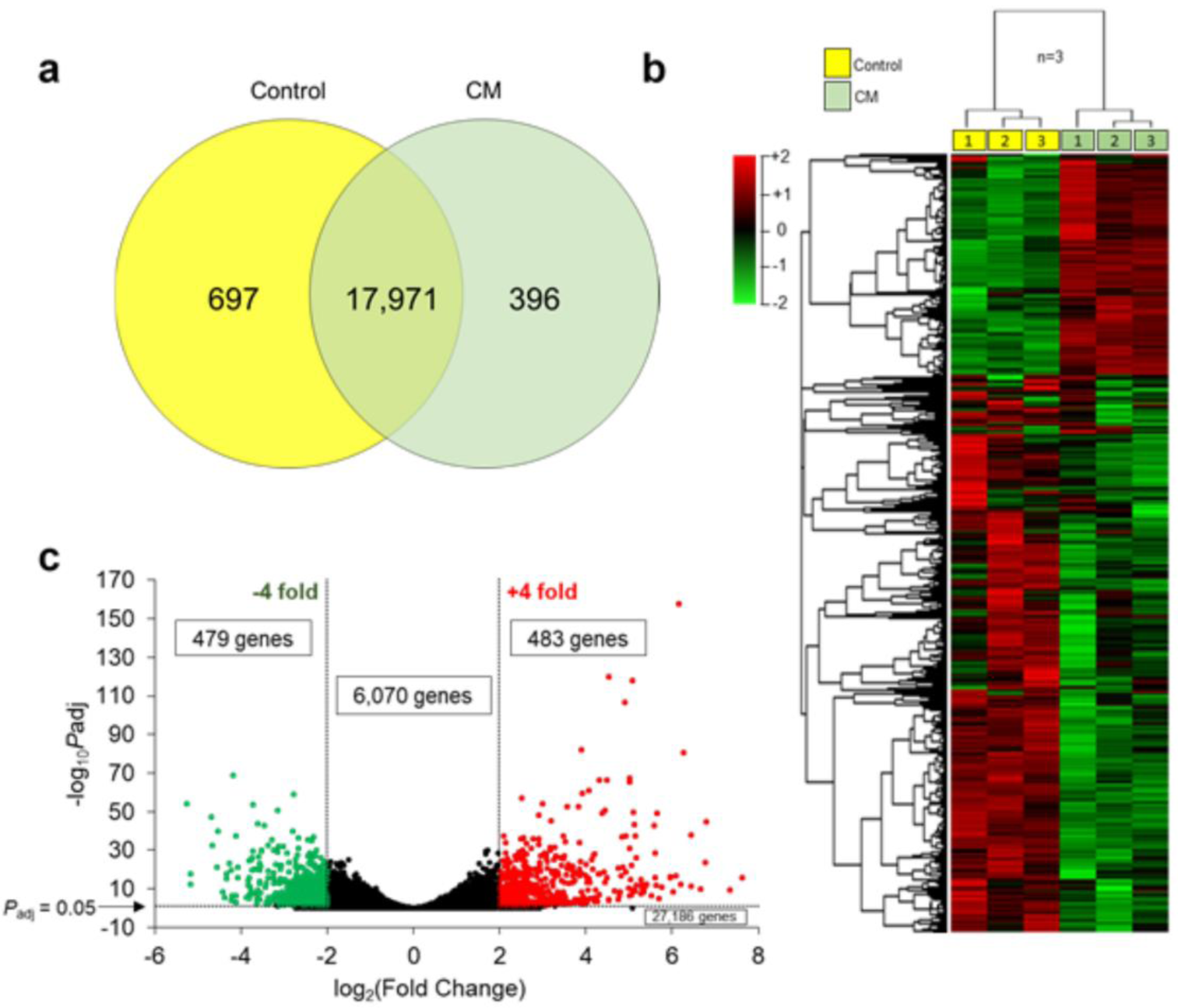
Transcriptomic changes in *A. thaliana* suspension cell cultures induced by *N. punctiforme*-CM. Changes in gene expression following a 1-hr incubation with either fresh *N. punctiforme* growth medium (Control) or *N. punctiforme*-CM (20% v/v) are shown. **a**. Venn diagram depicting co-expressed and uniquely expressed genes between control and CM treatments. **b**. Hierarchical clustering of 7,032 genes differentially expressed between control and CM treatments. **c**. Volcano plot depicting the distribution of all 34,218 annotated reads, where a 4-fold change cut-off in expression is delimited.

### Functional categorisation of genes regulated by *N. punctiforme*-CM

To functionally interpret these transcriptomic changes, we performed gene ontology (GO) and pathway enrichment analyses to systematically evaluate the biological significance of the genes which were 4-fold differentially expressed (Fig. 3). Up-regulated genes were mainly associated with defence responses, chitinase activity, indole-containing metabolic processes, and responses to chitin. Notable up-regulated pathways included phenylpropanoid biosynthesis, photosynthesis and glutathione metabolism. The most enriched processes associated with the down-regulated expression gene set were primary metabolic processes, such as transcription and the synthesis of macromolecules, responses to hormones, and meristem and shoot system development (Fig. 3). Photosynthesis was also suggested to be a down-regulated pathway, however, the expression of only 3 photosynthesis-associated genes were down-regulated, whereas the expression of 22 photosynthesis-associated genes were up-regulated (Supplementary Fig. 1), suggesting an overall up-regulation of photosynthesis. In general, the expression of hormone signalling-associated genes was down-regulated (Supplementary Fig. 1).

**Fig. 3.**
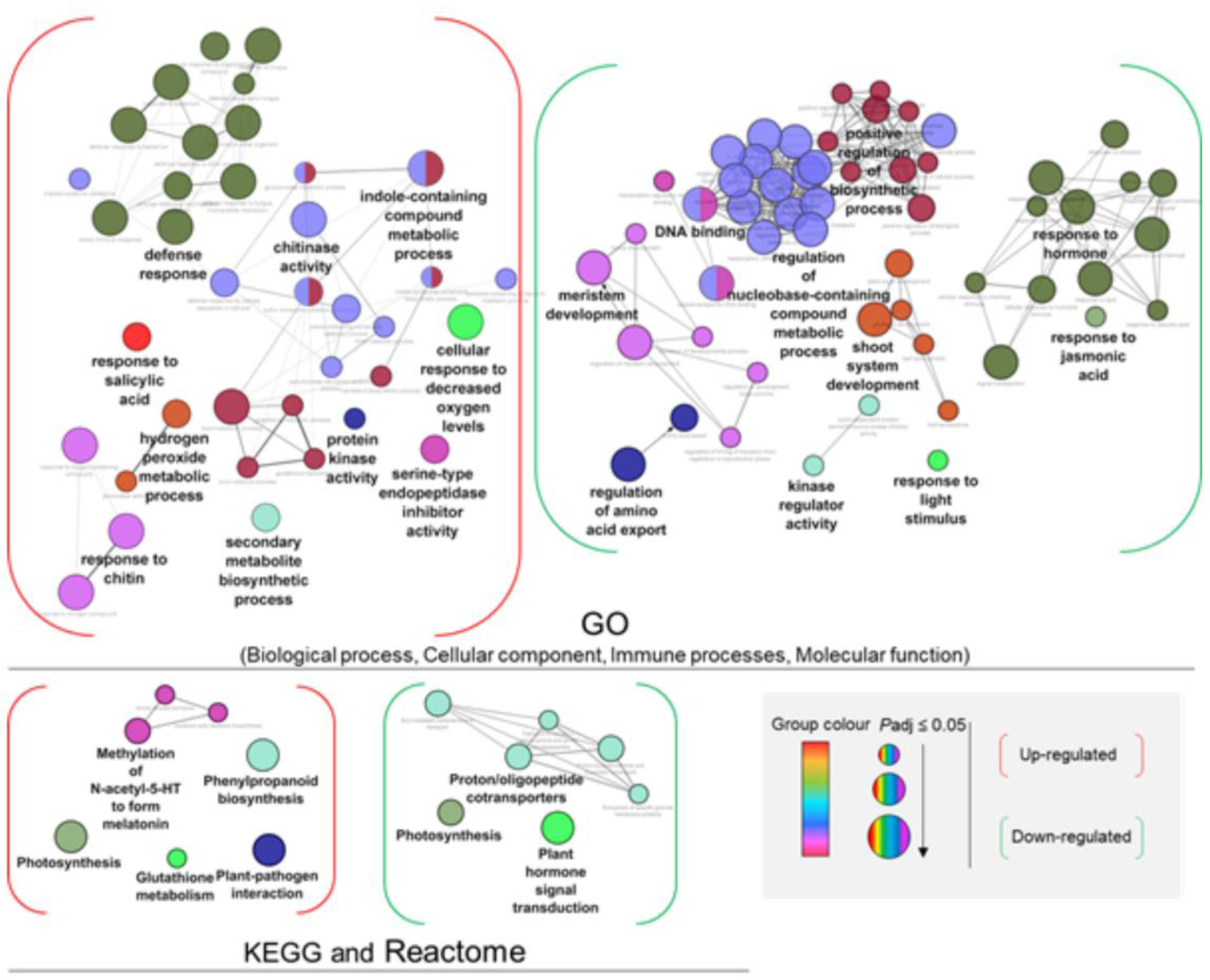
Functional categorisation of genes that were 4-fold differentially regulated in response to N. punctiforme-CM. GO terms and pathways (nodes) are connected by common genes. Terms in bold are the most statistically significant with respect to other terms in a group. As indicated in the legend, node colour corresponds to GO/ KEGG and Reactome group, whereas node size denotes statistical significance. The number of genes for each term are provided in Supplementary Fig. 1.

### Transcription factor (TF) enrichment analysis

To better understand the significance of the CM-induced transcriptional reprogramming, we analysed the TF composition of the differentially expressed gene set. Of the 962 differentially regulated genes, 121 were predicted to encode TFs (Supplementary Data 2). Expression of 24 TF genes belonging to 8 different families were up-regulated, with the most represented encoding WRKY family TFs (Supplementary Data 2). Conversely, a total of 97 TF genes spread across 27 families were significantly down-regulated, with genes encoding DOF and MYB family TFs being the most enriched (Supplementary Data 2). To investigate this further, we screened the promoters of the differentially expressed genes for TF-binding motifs and observed a significant enrichment of *cis*-motifs corresponding to known binding sites of 43 different TFs in the up-regulated gene set (462 genes) (Fig. 4). Of these, 41 were W-box recognition elements for WRKY TFs, with WRKY50 motifs being associated with 56 different genes. Only 9 TFs were associated with the down-regulated gene set, most of which belonged to the TCP class, a family of plant-specific developmental regulators (Li, 2015) (Fig. 4). A list of all the TF binding motifs can be found in Supplementary Data 3.

**Fig. 4.**
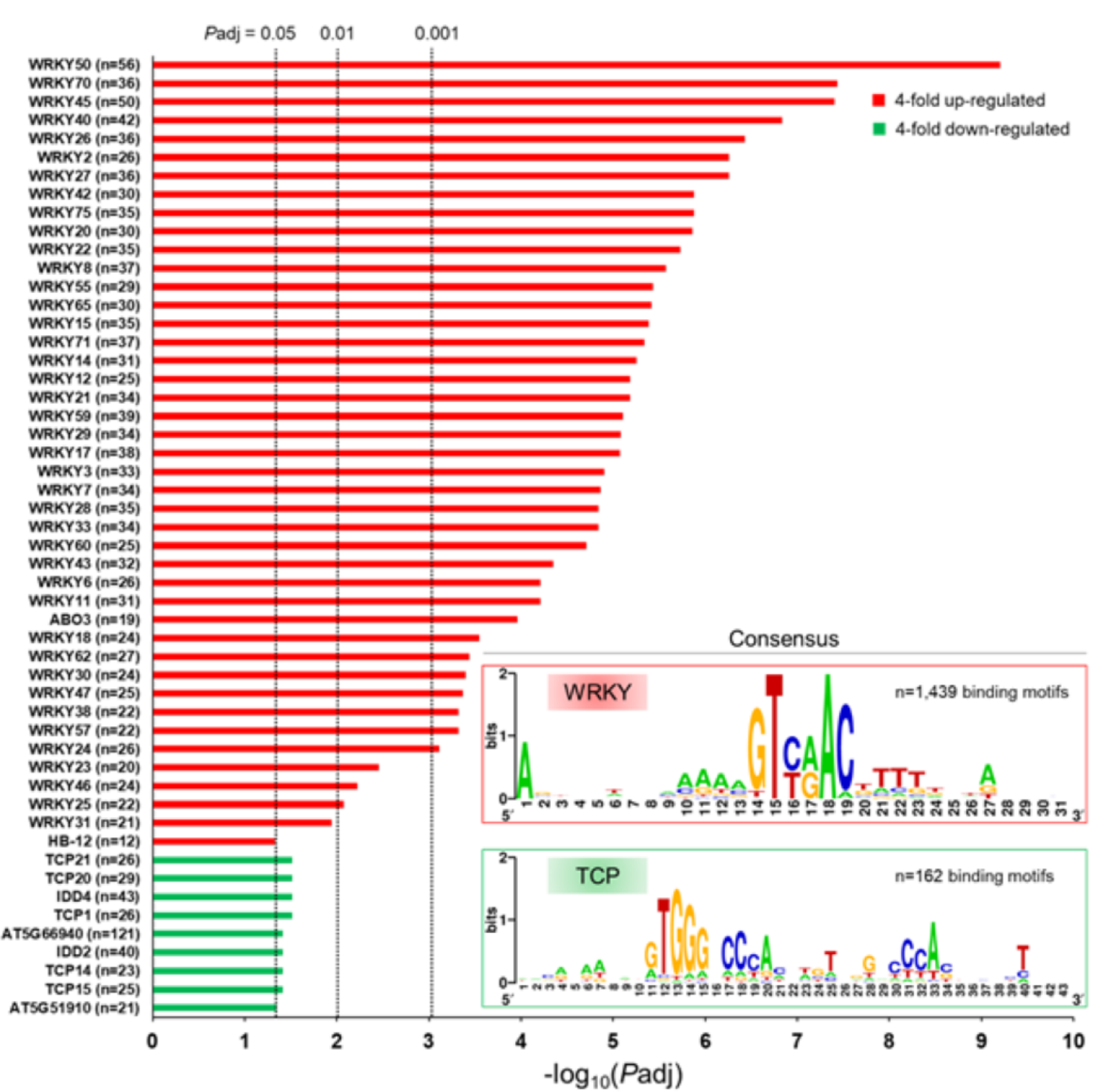
cis-Motif analysis of differentially expressed genes. Transcription factors (TFs) for which binding motifs were found in the promoter regions of the 962 differentially expressed genes (≥ 4-fold) in response to N. punctiforme-CM. For each TF, the number of genes for which canonical binding motifs were found is given. Also depicted are the enriched consensus sequences for WRKY and TCP family transcription factors.

### Analysis of potential protein-protein interaction networks

We generated a STRING protein-protein interaction network using the 962 significantly regulated genes to render the transcriptomic changes into an interaction network (Fig. 5). Of these, the STRING algorithm computed the protein products of 170 as being highly likely to interact with the product of at least one other differentially regulated gene. An additional 72 proteins were also linked in this network from the STRING database. The entire network comprised 1,752 edges (interactions), 103 hubs (proteins with 3 or more edges) and 40 components, the largest of which comprised 77 proteins (nodes), 45 of which were up-regulated (Supplementary Data 4). Most of these are implicated in defence responses to biotic and abiotic stresses, with WRKY33 appearing as a major hub linked with 20 different proteins. Also included were a range of proteins (*i*.*e*., IGMT1-3, BGLU26, CYP81F2 and CYP71A12) which are implicated in the metabolism of defence metabolites, such as glucosinolates and cyanogenic glycosides (Clay et al., 2009; Pfalz et al., 2016) (Fig. 5).

**Fig. 5.**
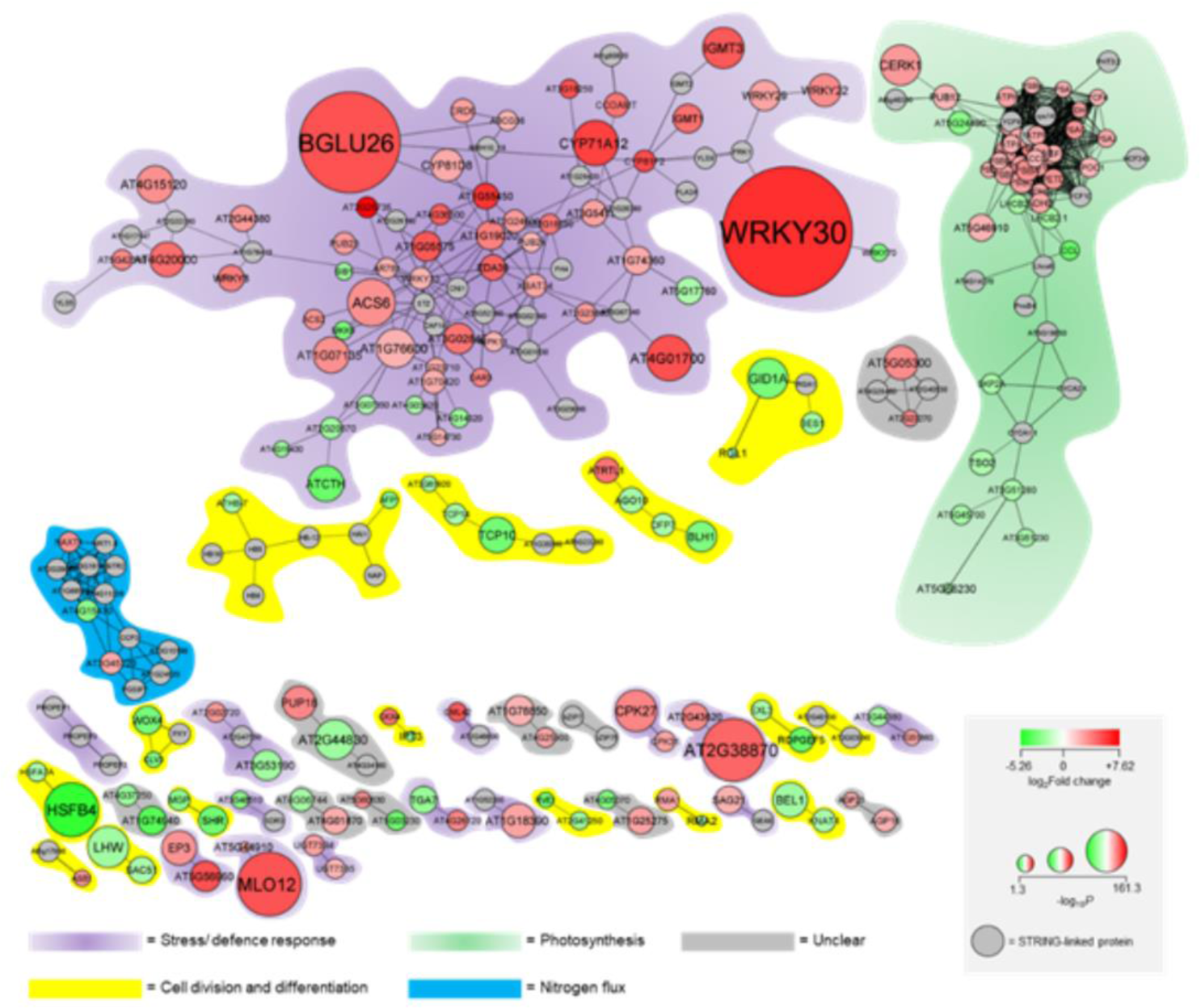
STRING protein-protein interaction network generated using genes that were at least 4-fold differentially regulated by *N. punctiforme*-CM. The network comprises 40 separate components which have been grouped into four different biological processes based on the predicted function of the majority of nodes (proteins) within each. As indicated in the legend, node colour and colour intensity correspond to change in gene expression level, node size denotes statistical significance of differential expression and grey nodes are associated through the STRING database. Network topology measurements can be found in Supplementary Data 4.

The next two largest components (58 and 12 nodes) mainly consisted of proteins which are associated with photosynthesis and the transport of nitrate and nitrite, respectively. The remaining 37 components (each comprising 8 nodes or less) appeared to be relevant in cell division and differentiation (15 components), stress/defence responses (12 components), or have functions that could not be easily curated (8 components) (Fig. 5). Included in the cell division and differentiation components are enzymes involved in cytokinin and brassinosteroid metabolism, as well as a range of proteins implicated in the regulation of cell fate. One such component comprises WOX4, CLV3 and PXY. *WOX4* and *CLV3*, which negatively regulate shoot stem cell identity (Somssich et al., 2016; Wahl et al., 2010), were strongly down-regulated (36- and 10-fold, respectively), suggesting that *N. punctiforme* may also negatively regulate cell differentiation.

## Discussion

### Is the regulation of PCD a requisite for the evolution of plant-cyanobacteria symbioses?

Symbiotic associations between microbes and higher eukaryotic organisms can be described in terms of the microbial symbiont evolving to tolerate or even suppress the innate immune responses of the eukaryotic host (Scwartzman and Ruby,2016). The symbiosomal environment of *Gunnera* spp., in which *N. punctiforme* can be found as intracellular cyanobionts, is characterised by a low pH and a high level of lysosomal enzymatic activity (Johansson and Bergman, 1992), suggesting that *Nostoc* cyanobionts possess the requisite traits to tolerate such conditions. It was recently demonstrated that *Nostoc* cyanobionts possessed 32 additional gene families compared to a free-living strain, and that intracellular plant-symbiotic Nostocales spp. likely evolved from a single common ancestor, whereas extracellular plant-symbiotic species likely evolved convergently (Warshan et al., 2017, 2018). This suggests that *Nostoc* cyanobionts may possess specialised suites of genes which immunologically permit symbiotic association. Importantly, we showed that conditioned medium (CM) from *N. punctiforme* did not trigger PCD but can actively attenuate stress-induced PCD (Fig. 1b). In a diverse range of bacteria-eukaryote symbioses, bacterial symbionts are known to produce molecules (‘symbiosis signals’) which maintain cellular homeostasis and suppress host PCD (An et al, 2014; Kitchen and Weis, 2017; Pannebakker et al., 2007).

We hypothesised that *N. punctiforme* may secrete symbiosis signals that are capable of suppressing PCD in plant hosts. Mechanistically, it is unclear how such signals could attenuate PCD given that their identities have not been elucidated. Nevertheless, we were able to gain insights into the molecular dialogue between the cyanobiont and plant cells. Here, the expression of 6 genes predicted to encode glutathione S-transferase (GST) enzymes were significantly up-regulated in response to *N. punctiforme* CM (Supplementary Data 5). GSTs are ubiquitous enzymes which can conjugate the reduced form of glutathione to a diverse range of substrates for detoxification (Sheehan et al., 2001). Many are inducible by a wide range of bacteria as part of the hypersensitive response (HR) in plants and they have been shown to reduce toxic ROS species and lipid hydroperoxides to limit the extent of HR-associated PCD^30^. It is tempting to hypothesise that *N. punctiforme* may induce the GST-mediated elimination of ROS and lipid hydroperoxides, thereby buffering against PCD activation. However, there are over 50 different *GST* genes in *A. thaliana* (Gullner et al., 2018), suggesting that only some of the GSTs up-regulated here are involved in regulating PCD. Importantly, this work is the first to demonstrate that a cyanobacterium is capable of suppressing plant PCD.

### *N. punctiforme* can modulate the expression of plant genes involved in photosynthesis, nitrogen flux, innate immunity, and cell growth

#### Photosynthesis and nitrogen flux

In total, 42 of the 962 differentially regulated genes mapped to the chloroplast, all of which were up-regulated and are mostly involved in photosynthesis (Supplementary Data 5). This included two genes encoding photosystem II reaction centre proteins (*psbJ* and *psbA*) and one encoding a photosystem I protein (*psaC*), which were up-regulated more than 8-fold. Plant chloroplasts possess functional signalling pathways which appear to derive from their free-living ancestors and are present in extant bacteria (Sugliani et al., 2016). It is tempting to suggest that some chloroplast regulatory elements are therefore responsive to bacterial MAMPs, particularly those of cyanobacterial origin (*i*.*e*., present within *N. punctiforme*-CM). The induction by cyanobionts of plant photosynthesis genes may also be an important symbiosis trait, as photosynthetic oxygen evolution is markedly down-regulated in the cyanobiont due to the preclusion of light in colonised tissues (Black and Osborne, 2004). To compensate, host photosynthesis rates likely must increase to facilitate the high energy requirement of nitrogen fixation in the cyanobiont. Indeed, the STRING-identification of a component harbouring proteins involved in nitrate and nitrite transport (*i*.*e*., NAXT1 and NRT1.8) could represent a physiological priming for the exchange of fixed carbon and nitrogen, which typifies all plant-cyanobacteria symbioses (Adams et al., 2012).

#### Innate immunity

The most enriched biological processes from the up-regulated gene set were associated with plant defence and immunity. This was mirrored in the STRING network, in which the largest component primarily comprised of up-regulated genes encoding proteins involved in stress and defence responses (Fig. 5). The induction of defence genes would presumably have been preceded by the recognition of *N. punctiforme* symbiosis signals, although their identities and their cognate plant receptor(s) are at present unclear. A total of 17 receptor kinases were significantly up-regulated (Supplementary Data 5), including *FLS2* and *CERK1*, both of which were induced 7.5-fold. FLS2 and CERK1 are involved in the perception of the Flg22 peptide and chitin MAMPs, respectively, which are transduced into the transcription of defence genes via mitogen-activated protein kinase (MAPK) signalling cascades (Sheikh et al., 2016; Yamada et al., 2016). Although the enrichment analysis suggested that responses to chitin were up-regulated, *N. punctiforme* does not produce chitin (nor flagellin). However, N-acetylglucosamine, the amino sugar from which chitin oligomers are synthesised, is a common constituent of cyanobacterial exopolysaccharides and is produced by Nostocales species (Halsør et al., 2019; Rossi et al., 2015). Interestingly, CERK1 homologues in legumes can recognise Nod factors from symbiotic Rhizobia (Zhang et al., 2015), but whether CERK1 in *A. thaliana* can also perceive symbiosis signals from *N. punctiforme* remains to be determined. However, the hypothesis that extracellular amino sugars are important as symbiosis signals in *N. punctiforme* remains speculative. The few studies which have investigated the composition of the exo-proteome and exo-metabolome of symbiotic *Nostoc* spp. have revealed an enrichment of small peptides, polyketides and peptidoglycan-derived tetrasaccharides (Liaimer et al., 2016; Vilhauer et al., 2014). At present, other than a group of polyketides which were demonstrated to be important in suppressing the motile growth phase of *N. punctiforme* during symbiotic association (Liaimer et al., 2015), none of these compounds have been functionally characterised.

Once putative symbiosis signals from *N. punctiforme* are perceived, presumably, they may be transduced through MAPK cascades into downstream transcriptional reprogramming. In total, the expression of 3 genes predicted to encode MAPKs (1 MAPK and 2 MAPKKKs) were significantly up-regulated, whereas 2 were significantly down-regulated (1 MAPKK and 1 MAPKKK; Supplementary Data 5). A strikingly wide range of WRKY TFs were associated with the up-regulated gene set, which potentially could be downstream targets of any MAPK signalling cascade (Jiang et al., 2017). WRKY TFs are master regulators of biotic stress responses (Phukan et al., 2016) and their W-box binding motifs were highly represented in the promoter regions of the up-regulated gene set (Fig. 5). Eight different WRKY TFs were themselves strongly up-regulated (Supplementary Table 2), indicating that they may be critical in coordinating plant responses to symbiotic cyanobacteria. Although few plant genes which are regulated by cyanobionts have been elucidated, a recent study showed that the symbiotic association between the water fern, *Azolla filiculoides* and its cyanobiont, *Nostoc azollae* resulted in the positive and negative regulation of 88 and 72 genes, respectively (Eily et al., 2019). These genes were termed “putative symbiosis genes” and were selected on the basis of their differential expression in the *A. filiculoides* host in the presence of *N. azollae* but in the absence of a source of combined nitrogen, which was reasoned to be the “optimal symbiosis growth condition”. The *Nostoc*-*Azolla* system (Eily et al., 2019) obviously differed radically to the one used here, in which nitrogen-supplemented, un-differentiated *A. thaliana* cells were exposed to CM derived from vegetative *N. punctiforme* cell cultures (non-diazotrophic). Nonetheless, both approaches identified a range of WRKY TF genes as being differentially regulated by *Nostoc* cyanobionts, including *WRKY72*, which was strongly up-regulated in both systems.

WRKY72, an inducible TF which activates a defence transcriptome that is largely independent of the defence hormone, salicylic acid (SA) (Bhattarai et al., 2010) was induced 30-fold by *N. punctiforme* in *A. thaliana*. In contrast, *WRKY30*, which was one of the most strongly up-regulated genes (72-fold), is SA-inducible and its over-expression in rice results in the accumulation of another defence hormone, jasmonic acid (JA), as well as JA-inducible genes (Peng et al., 2012). SA and JA responses were GO terms associated with the up- and down-regulated gene sets, respectively, whereas “hormonal signalling” was associated only with the latter (Fig. 3). It is however worth cautioning that the roles of both hormones in regulating plant-bacteria symbioses are not easily generalised. For instance, whether SA and JA signalling, which are antagonistic within individual plants (Van der Does et al., 2013), positively or negatively regulate Rhizobia-induced nodulation appears to be dependent on the legume host species (Liu et al., 2018).

It is noteworthy that the strong induction of immunity-associated genes was not followed by an incompatible response (*i*.*e*., PCD). Rather, cells were primed against the activation of PCD. That this occurred in the non-host *A. thaliana* may not be that surprising given the observation that in both *A. thaliana* and host legume spp., Rhizobia initially activate immune responses (ROS accumulation, cytosolic Ca^2+^ burst, MAPK signalling) only for Nod factors to quickly suppress them. Nod factors can also induce the down-regulation of receptor kinases, such as FLS2^7^. Here, although FLS2 was up-regulated, 10 different receptor kinases were down-regulated by *N. punctiforme*-CM (Supplementary Data 5), which may signify a suppression of immunological receptivity. It is also worth noting that *N. punctiforme* does not possess the type III secretory systems associated with the delivery of effector proteins by many pathogenic and some symbiotic bacteria (Schuergers and Wilde, 2015). This would indicate that either the perception of symbiosis signals (*i*.*e*., MAMPs) is by itself sufficient to suppress host immune responses, or signalling molecules analogous to effectors can be delivered without a secretory system. Nonetheless, the pattern of defence signalling glimpsed in this transcriptome may be viewed as an immunological signature which, to varying degrees, could be conserved in plant hosts upon acquisition of their cyanobionts.

#### Cell growth

Whilst a distinctive defence transcriptome was strongly induced, the expression of genes which were down-regulated by *N. punctiforme* appeared to primarily be important in regulating cell division and differentiation. The main down-regulated processes were associated with primary metabolic activity, such as the metabolism of nucleic acids, the biosynthesis of cyclic and aromatic compounds (presumably nucleobases) and transcription (Fig. 3). This may be explicable given that many of the down-regulated genes are associated with regulating cell division and meristem development. For instance, *WOX4, CLV3* and *FAF4* were all strongly down-regulated (36-, 10- and 9-fold, respectively). WUSCHEL-related HOMEOBOX 4 (WOX4) positively regulates procambium differentiation (Etchells et al., 2013), whereas CLV3 and FAF4 negatively regulate WUSCHEL (WUS), which is a meristem-specific transcription factor that positively regulates shoot stem cell identity (Somssich et al., 2016; Wahl et al. 2010). Although WUS itself was not expressed, likely because the cell cultures are not meristematic, the down-regulation of *CLV3, FAF4* and *WOX4* together suggests that *N. punctiforme* can negatively regulate cell differentiation.

Genes involved in the regulation of cell division were also strongly down-regulated; *MYB59*, which is a negative regulator of cell proliferation (Mu et al., 2009), was down-regulated 21-fold. MYBs were the second-most represented TFs in the down-regulated gene set, with DOF family TFs being the most abundant. DOF and MYB TFs are involved in a range of developmental processes (Ambawat et al., 2013; Le Hir and Bellini, 2013), which suggests that broadly, defence signalling was up-regulated at the expense of developmental signalling. At least 5 genes involved in auxin biosynthesis were significantly down-regulated (Supplementary Data 5), whereas *LOG1*, which encodes a cytokinin riboside 5’-monophosphate phosphoribohydrolase that converts cytokinin into its active form (Kuroha et al. 2009), was strongly up-regulated (5.7-fold). The regulation of hormones associated with the control of cell growth and division may be of significance. Nodule formation in legumes is preceded by the induction of de-differentiation and cell division in root cortical cells, and the induction by *N. punctiforme* of multiple rounds of cell division within the glands of *Gunnera* spp. is expected to be critical for symbiosis establishment (Geurts et al., 2016). Potentially, the control of cell division and differentiation by plant-symbiotic cyanobacteria is a rudimentary trait which is reflected in the down-regulated transcriptome here.

Based on our data presented in this study, we propose that symbiosis signals from symbiotically-competent cyanobacterial cells are recognised by plant receptor kinases. Upon signal perception, a signal transduction cascade potentially involving MAPK intermediates activates a range of WRKY TFs. WRKY TF hubs may drive the expression of multiple defence-associated genes, including those encoding GST enzymes. This induction could be critical in buffering cells against the activation of PCD (Fig. 6), thereby immunologically favouring symbiotic association.

**Fig. 6.**
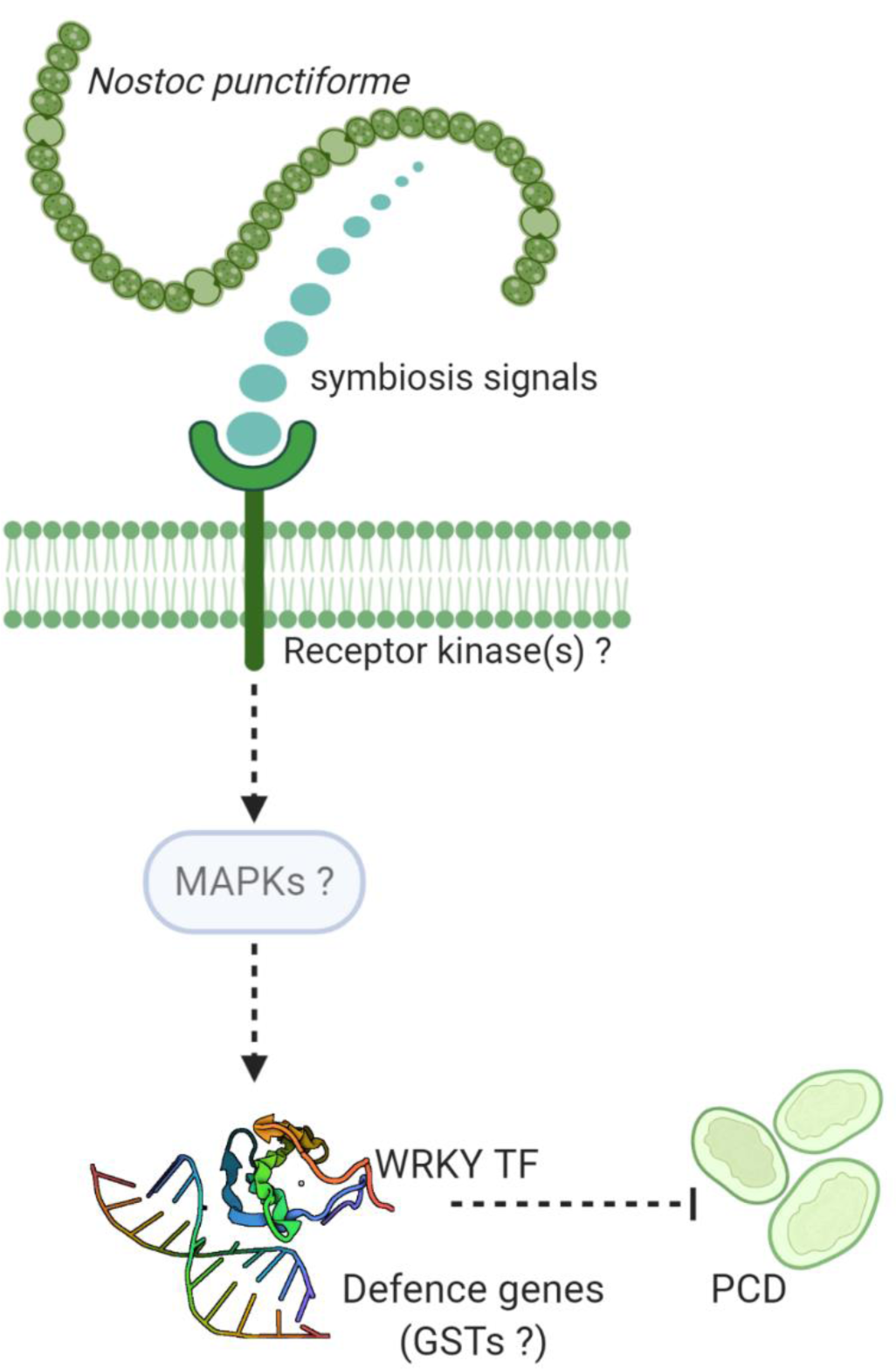
Proposed model by which symbiotic Nostoc spp. may affect plant host cell immunity. Cyanobiont-derived symbiosis signals binds a plant receptor kinase(s) followed by signal transduction, potentially through a MAPK signalling cascade, into the activation of WRKY family TFs. WRKYs may drive the expression of a range of defence genes, including some which encode GST enzymes. Downstream of this, host cells could be primed against the induction of PCD.

In summary, our results demonstrate for the first time that the cyanobiont, *N. punctiforme* is capable of suppressing plant PCD, which is a hallmark immune response to incompatible microbes. This response appears to be preceded by the up-regulation of a range of immunity-associated genes, such as those encoding WRKY TFs and GST enzymes. In addition, a range of genes which function in the regulation of cell division and differentiation were down-regulated. Our study brings to light the early responses of plant cells to cyanobionts, and the molecular determinants that are activated to attenuate PCD. Furthermore, our study provides the molecular blueprint for future endeavours to better understand this aspect of these enigmatic cross-species relationships.

## Methods

### Growth of *N. punctiforme* cells and preparation of conditioned medium

*N. punctiforme* strain ATCC 29133 was grown in liquid BG-11 (NH_4_) medium at 21°C with shaking at 120 rpm for a photoperiod of 16 hrs light (*ca*. 165 μmol m^-2^ sec^-1^ of cool white fluorescent light) and 8 hrs darkness (Campbell et al., 2007; Christman et al., 2011). To harvest CM, a volume of cells equivalent to 60 μg of chlorophyll *a* was added to 100 ml of fresh growth medium. After 21 days of growth (stationary phase), cells were harvested by centrifugation at room temperature for 20 min at 3,000 *g* in glass Corex™ round-bottomed tubes. The supernatant was then passed through two pieces of Whatman™ grade I filter paper and collected in chromic acid-washed glass vials before storing at -20°C for subsequent use.

### Maintenance of *A. thaliana* suspension cell cultures

*A. thaliana* var. Ler-0 suspension cell cultures were maintained under the same growth conditions as above but in Murashige and Skoog (MS) medium (Murashige and Skoog, 1962) containing 3% sucrose, 0.5 mg L^-1^ naphthaleneacetic acid (NAA) and 0.05 mg L^-1^ kinetin (pH adjusted to 5.8 using KOH). Cells were sub-cultured every 7 days by transferring 10 ml of mature cells into 90 ml of fresh growth medium.

### Monitoring the effects of *N. punctiforme*-CM on PCD in *A. thaliana* cells

To study the effects of *N. punctiforme*-CM on PCD in *A. thaliana* cells, 2.5 ml of CM was added to 10 ml of a 7-day-old *A. thaliana* suspension cell culture (cell density = ca. 5 × 10^4^ cells ml^-1^) and left to pre-incubate for 1 hr under standard culturing conditions. Next, cells were either incubated at room temperature (∼21°C) or placed in a water bath set to 51°C for 10 min with constant agitation at 120 rpm (Kacprzyk et al., 2017). After returning to standard culturing conditions for 24 hr, the frequency of viable, PCD-committed and necrotic cells was measured. This was done by mixing cells in a 1:1 volume of 0.004% fluorescein diacetate (FDA; w/v in 100% acetone) before quantification using epifluorescence microscopy (excitation wavelength = 490 nm, emission wavelength = 515 nm) (Alden et al., 2011). Green fluorescence as a result of FDA hydrolysis was indicative of cell viability. A retracted protoplasm and loss of green fluorescence indicated PCD, whereas a lack of fluorescence and no retracted protoplasm was indicative of cell necrosis (Reape et al., 2008). To determine statistical significance, arcsine-transformed proportions were subjected to a two-tailed Student’s t-test (*P* ≤ 0.05).

### RNA isolation and quality control

RNA was isolated from flash-frozen 1 ml aliquots of *A. thaliana* cells immediately prior to PCD induction. Frozen cells were disrupted by bead beating at 30 h/z for 1 min before purifying total RNA using an RNeasy^®^ plant mini kit (Qiagen™; ref. 74904) according to manufacturer’s instructions. RNA was quantified using a NanoDrop^®^ 1000 spectrophotometer and quality determined on a 1% TBE agarose gel. A Nano 6000 Assay Kit was used with a 2100 BioAnalyzer^®^ system (Agilent^©^) revealed that all sample RIN scores were over 9.0 (Supplementary Fig. 2).

### cDNA library preparation

Library preparation and RNA-seq was performed by Novogene (Beijing, China). A total of 1 μg RNA was used as input material for library generation using the NEBNext^®^ UltraTM RNA Library Prep Kit for Illumina^®^ (NEB USA), at which point index codes were added to each sample. mRNA was purified using magnetic beads harbouring poly-dT oligos. Fragmentation was performed using divalent cations in heated NEBNext First StrandSynthesis Reaction Buffer. First strand cDNA was synthesised using random hexamers and M-MuL MuLV Reverse Transcriptase. Second strand cDNA was synthesised using DNA Polymerase I (+RNase H). Blunt ends were generated via exonuclease/ polymerase activities. Next, 3’ DNA ends were adenylated before ligating an NEBNext hairpin loop adaptor for hybridization. Fragments of *ca*. 200 bp were preferentially selected for using an AMPure XP system (Beckman Coulter, Beverly, USA). Purified, adaptor-ligated fragments were treated with 3 µl of USER^®^ Enzyme (NEB, USA) before selective PCR enrichment using Phusion^®^ High-Fidelity DNA polymerase using Universal and index primers. Products were purified again using the AMPure XP system and library quality assessed using the 2100 BioAnalyzer^®^ system.

### Sequencing and transcriptome assembly

Index-coded samples were clustered on a cBot Cluster Generation system using a PE Cluster Kit cBot-HS (Illumina) followed by paired-end sequencing on an Illumina NovaSeq 6000 platform. FASTAQ raw reads were filtered to remove adaptors, poly-N sequences and low-quality reads. Q20, Q30 and GC content were also calculated (Supplementary Data 6). Using HISAT2 software, cleaned-up reads were mapped to the *A. thaliana* genome (TAIR10) before expressing read quantification as FPKM.

### Differential expression analysis

Differential expression of genes between treatments (n = 3 biological replicates) was determined using the DESeq2 R package. The false discovery rate (FDR) was determined using the Benjamini and Hochberg procedure, after which genes with an adjusted *P* value (*P*adj) of 0.05 were considered to be differentially expressed. Hierarchical clustering of the log_10_FPKM values of differentially expressed genes was performed using the gplot R package.

### Enrichment analysis

Gene ontology (GO) and pathway enrichment analyses were performed on the 4-fold differentially regulated gene set. This was done using the ClueGO plugin (V2.5.7) (Bindea et al., 2009) for Cytoscape (V3.7.2) (Shannon et al, 2003), with the following settings: Kappa Score threshold = 0.5, Min GO level = 3, Max GO level = 8, Min genes per cluster = 3, Min % genes = 4, GO Term Fusion = True. The cut-off for enriched terms was set to a familywise error rate of *P*adj ≤ 0.05.

### TF enrichment analysis

Peptide sequences belonging to the 962 differentially expressed genes were acquired in FASTA format using BioMart (Smedley et al., 2015) (version 0.7) through the EnsemblPlants genome resource portal (https://plants.ensembl.org/biomart/martview/5b18fb7d060dfa269fe3dadd41bfbc2a). The TF composition of this data set was ascertained using the TF prediction tool at the plant TF database PlantTFDB (Jin et al., 2017) (http://planttfdb.cbi.pku.edu.cn/prediction.php). The promoter regions (region spanning 600 bp upstream of the 5’-UTR) of the 962 differentially expressed genes were also extracted using BioMart. TF names for which *cis*-binding motifs were enriched in these promoter sequences were acquired using the regulation prediction tool available through the PlantRegMap (Tian et al., 2020) portal (http://plantregmap.cbi.pku.edu.cn/regulation_prediction.php). The FDR was controlled using a Benjamini-Hochberg correction (*P*adj ≤ 0.05).

### STRING network analysis

The STRING protein-protein interaction network was generated based on the 4-fold differentially regulated gene set using the stringApp plugin (V1.5.1) (Doncheva et al., 2019) for Cytoscape. All possible evidence channels were selected and a confidence score of 0.7 (high strength) was selected (*P*adj ≤ 0.05).

## Acknowledgements

This work was financially supported by a Government of Ireland Postgraduate Scholarship from the Irish Research Council (GOIPG/2015/2695) to SPB.

