## Supplementary Data 2 for "The nitrogen-fixing symbiotic cyanobacterium, *Nostoc punctiforme* can regulate plant programmed cell death"

**Supplementary Data 2** List of transcription factors (TFs) extracted from the 962 genes that were differentially regulated by *N. punctiforme*-coditioned medium (CM). Red = up-regulated, green = down-regulated.

| TF Family | Individual TFs |
|---|---|
| **Up-regulated (n=24)** | |
| WRKY | WRKY8, WRKY22, WRKY28, WRKY29, WRKY30, WRKY33, WRKY41, WRKY72 |
| C2H2 | AT3G53600, STZ, ZAT11, ZF1 |
| MYB | MYB2, MYB49, MYB87, MYB120 |
| NAC | AT3G12910, NAC042, NAC090 |
| ERF | AT1G71520, AT2G33710 |
| bHLH | AT5G56960 |
| GRF | GRF8 |
| LBD | LBD32 |
| **Down-regulated (n=97)** | |
| DOF | AT1G64620, AT1G69570, AT2G28810, AT2G34140, AT5G02460, CDF1, HCA2, OBP4, TMO6 |
| MYB | AT5G05790, MYB13, MYB27, MYB34, MYB59, MYB61, MYB100, MYB107 |
| ERF | AT2G44940, AT5G07580, AT5G61590, ESE3, ESR1, Rap2.6L, SHN3 |
| HD-ZIP | ATHB13, HAT3, HB5, HB6, HB7, HB12, HB16, |
| NAC | NAC011, NAC028, NAC047, NAC074, NAC3, NAM, NAP |
| bHLH | AT2G40200, bHLH071, bHLH093, FBH4, MYC4, PIL1 |
| bZIP | bZIP24, bZIP44, bZIP7, bZIP75, DPBF2, TGA7 |
| C2H2 | AT2G29660, AT5G03510, IDD12, IDD14, MGP, NUC |
| MYB_related | AT1G70000, AT5G58900, EPR1, RVE1, TRFL2 |
| TCP | AT1G35560, AT2G37000, AT5G23280, TCP10, TCP14 |
| GRAS | AT2G37650, RGA1, RGL1, SHR |
| TALE | BEL1, BLH1, BLH3, KNAT4 |
| AP2 | ANT, RAP2.7, TOE3 |
| G2-like | AT3G25790, AT4G37180, MYBC1 |
| HSF | HSFA7A, HSFB4 |
| Trihelix | AT1G76870, GTL1 |
| WOX | WOX4, WOX7 |
| WRKY | WRKY35, WRKY70 |
| ARR-B | RR14 |
| BES1 | BZR1 |
| C3H | ATCTH |
| CO-like | BBX15 |
| DBB | STH |
| GRF | GRF4 |
| LBD | LBD4 |
| M-type_MADS | AGL49 |
| NF-YB | NF-YB3 |
