## Supplementary Figures for "The nitrogen-fixing symbiotic cyanobacterium, *Nostoc punctiforme* can regulate plant programmed cell death"

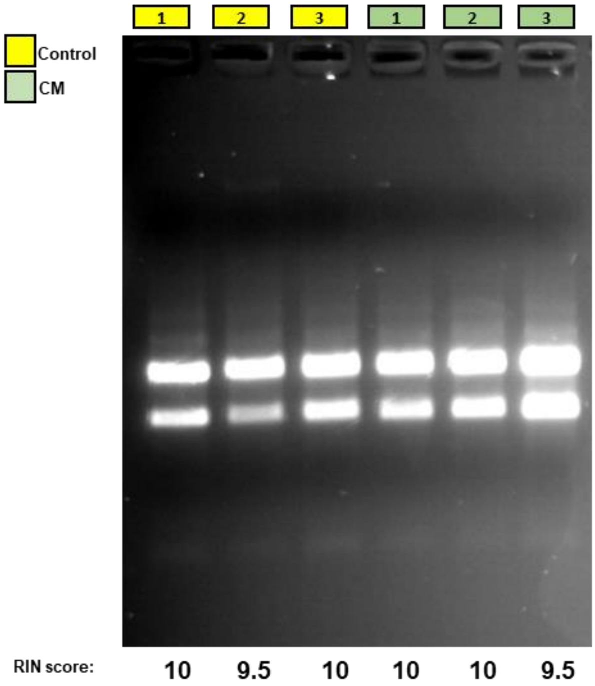


**Supplementary Fig. 1** Gel electrophoresis separation of total RNA samples isolated from *A. thaliana* cell suspension cultures using an RNeasy® plant mini kit (Qiagen™). Depicted are samples (n = 3) treated with fresh *N. punctiforme* growth medium (Control) and *N. punctiforme*-conditioned medium (CM). Also indicated are sample RIN scores, as determine using a 2100 BioAnalyzer® system (Agilent©).

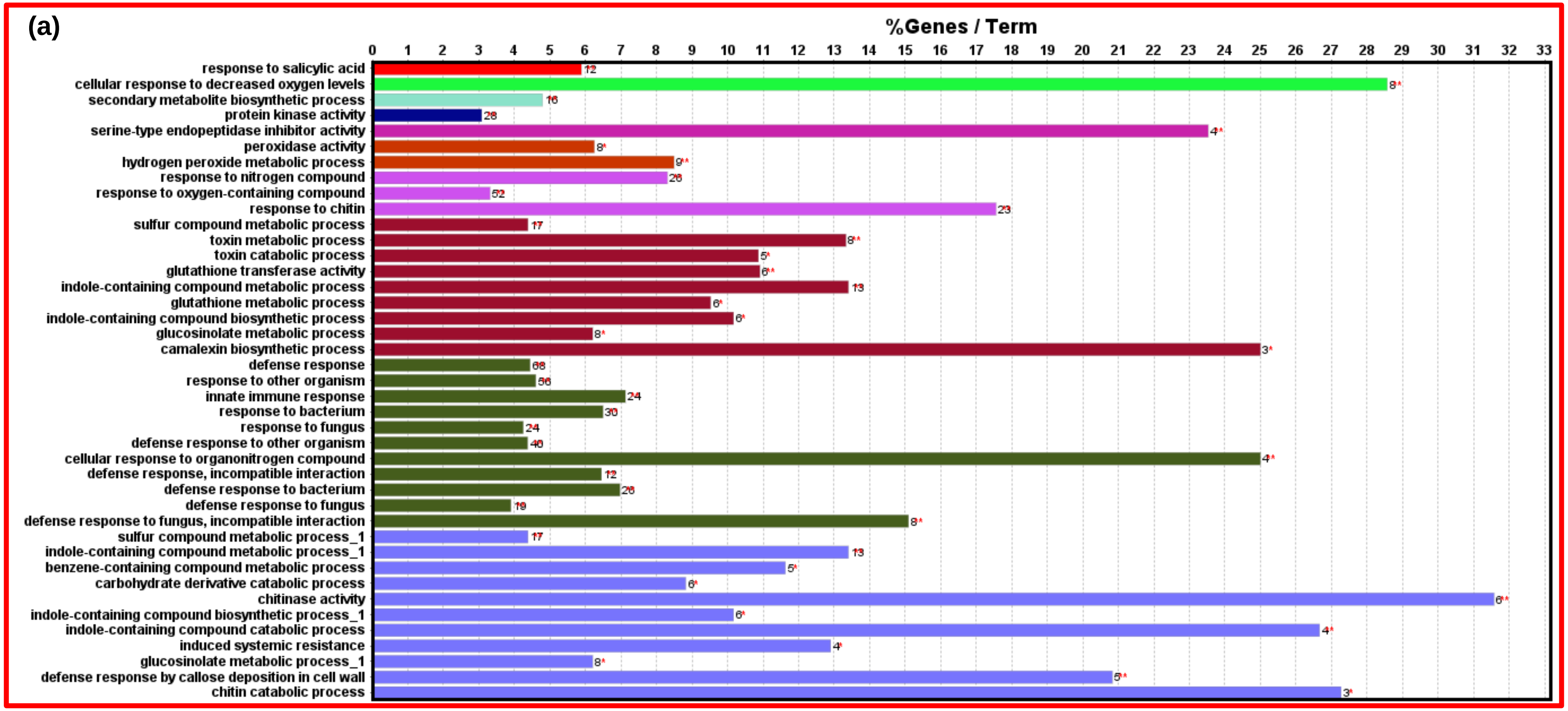
(a)
%Genes / Term
response to salicylic acid
cellular response to decreased oxygen levels
secondary metabolite biosynthetic process
protein kinase activity
serine-type endopeptidase inhibitor activity
peroxidase activity
hydrogen peroxide metabolic process
response to nitrogen compound
response to oxygen-containing compound
response to chitin
sulfur compound metabolic process
toxin metabolic process
toxin catabolic process
glutathione transferase activity
indole-containing compound metabolic process
glutathione metabolic process
indole-containing compound biosynthetic process
glucosinolate metabolic process
camalexin biosynthetic process
defense response
response to other organism
innate immune response
response to bacterium
response to fungus
defense response to other organism
cellular response to organonitrogen compound
defense response, incompatible interaction
defense response to bacterium
defense response to fungus
defense response to fungus, incompatible interaction
sulfur compound metabolic process_1
indole-containing compound metabolic process_1
benzene-containing compound metabolic process
carbohydrate derivative catabolic process
chitinase activity
indole-containing compound biosynthetic process_1
indole-containing compound catabolic process
induced systemic resistance
glucosinolate metabolic process_1
defense response by callose deposition in cell wall
chitin catabolic process

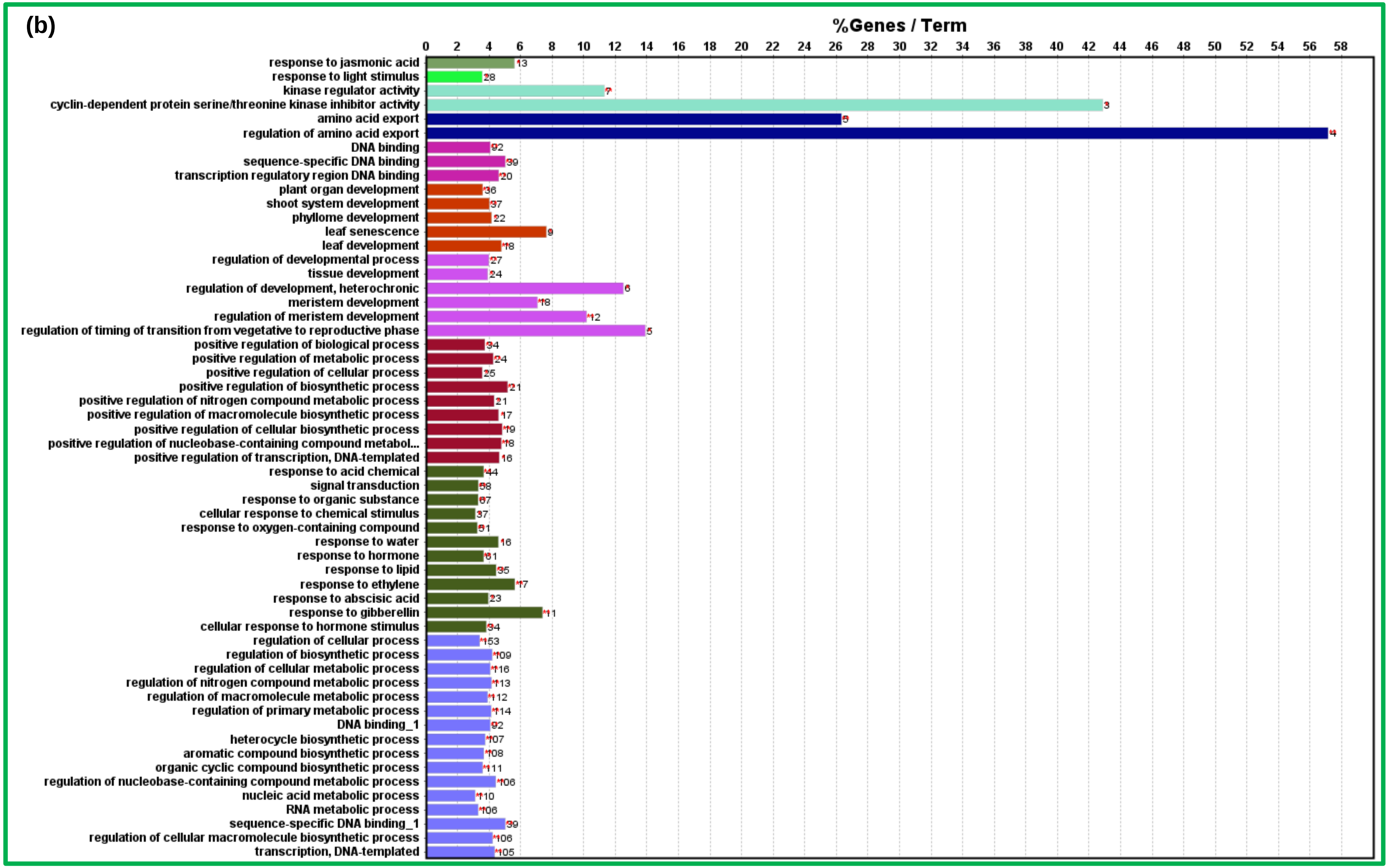
(b)
%Genes / Term
0 2 4 6 8 10 12 14 16 18 20 22 24 26 28 30 32 34 36 38 40 42 44 46 48 50 52 54 56 58
response to jasmonic acid 13
response to light stimulus 28
kinase regulator activity 7
cyclin-dependent protein serine/threonine kinase inhibitor activity 3
amino acid export 5
regulation of amino acid export 4
DNA binding 92
sequence-specific DNA binding 39
transcription regulatory region DNA binding 20
plant organ development 36
shoot system development 37
phyllome development 22
leaf senescence 9
leaf development 18
regulation of developmental process 27
tissue development 24
regulation of development, heterochronic 6
meristem development 18
regulation of meristem development 12
regulation of timing of transition from vegetative to reproductive phase 5
positive regulation of biological process 34
positive regulation of metabolic process 24
positive regulation of cellular process 25
positive regulation of biosynthetic process 21
positive regulation of nitrogen compound metabolic process 21
positive regulation of macromolecule biosynthetic process 17
positive regulation of cellular biosynthetic process 19
positive regulation of nucleobase-containing compound metabol... 18
positive regulation of transcription, DNA-templated 16
response to acid chemical 44
signal transduction 58
response to organic substance 67
cellular response to chemical stimulus 37
response to oxygen-containing compound 51
response to water 16
response to hormone 61
response to lipid 35
response to ethylene 17
response to abscisic acid 23
response to gibberellin 11
cellular response to hormone stimulus 34
regulation of cellular process 153
regulation of biosynthetic process 109
regulation of cellular metabolic process 116
regulation of nitrogen compound metabolic process 113
regulation of macromolecule metabolic process 112
regulation of primary metabolic process 114
DNA binding_1 92
heterocycle biosynthetic process 107
aromatic compound biosynthetic process 108
organic cyclic compound biosynthetic process 111
regulation of nucleobase-containing compound metabolic process 106
nucleic acid metabolic process 110
RNA metabolic process 106
sequence-specific DNA binding_1 39
regulation of cellular macromolecule biosynthetic process 106
transcription, DNA-templated 105

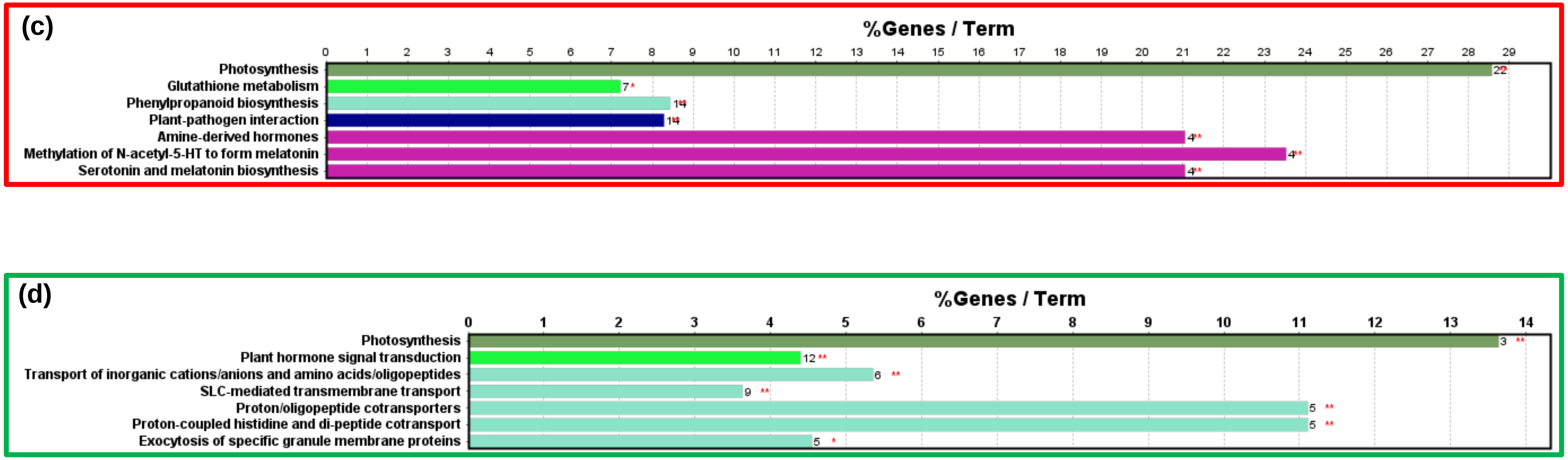


**Supplementary Fig. 2** Quantitative breakdown of GO and pathway enrichment analyses of *A. thaliana* genes which were differentially regulated by *N. punctiforme*-conditioned medium (CM). Shown are the number of genes linked to each term as well as the % coverage for all known genes for each term. Values are given for **(a)** up- and **(b)** down-regulated GO terms, as well as **(c)** up- and **(d)** down-regulated pathway terms.
